# Magnetic field enables spin matching in spin-selective pathways to enhance current in bacteria

**DOI:** 10.64898/2026.06.16.732578

**Authors:** Kshitij Kathait, Rahul Mishra, Lucinda Elizabeth Doyle

## Abstract

Electroactive bacteria export metabolic electrons through cytochrome-based extracellular electron transfer (EET) conduits. Although purified *Shewanella oneidensis* EET cytochromes are chiral and spin-selective, the functional impact of chirality-induced spin selectivity in live cultures remains unknown. Here, we show that a magnetic field enhances current in live *S. oneidensis*. EET to non-magnetic electrodes increased by ∼70 %, whereas electron uptake was insensitive to magnetic field. In contrast, electron uptake increased by >100 % when spin-polarised electrons were supplied from a magnetised electrode. This directional response suggests that magnetic field improves spin matching in the EET conduit, but only when the incoming electron flux is spin-polarised, implying that metabolic electrons are already spin-polarised before entering the conduit. Cytochromes in the conduit were downregulated despite higher current, indicating enhanced EET efficiency.

---

The passage of electrons through chiral molecules gives rise to chiral-induced spin selectivity (CISS) in which the molecular handedness couples with electron spin. Thus, chiral molecules act as spin filters and preferentially transmit electrons with one spin orientation (*1–5*). A range of biological molecules support such spin-selective electron transport when tested *in vitro* (*6–11*). However, whether molecular-scale spin selectivity can influence organism-level biological function remains largely unexplored.

Electroactive microorganisms provide a powerful platform to address this question because their metabolic electrons are exported across the cell envelope to an external electrode i.e. extracellular electron transfer (EET), enabling their real-time electrochemical analysis (*12–16*). This direct electrical access is not available in non-electroactive organisms, where metabolic electrons are retained within the cell or transferred to soluble electron acceptors. In the model electroactive bacterium *Shewanella oneidensis*, EET is mediated by iron-containing multiheme cytochromes that connect intracellular metabolism to solid electron acceptors (*17–23*). Notably, several cytochromes involved in *S. oneidensis* EET have been isolated and shown to exhibit spin-selective/filtering behaviour (*24*, *25*). Yet, whether this spin selectivity has a functional consequence during current generation in live *S. oneidensis* cultures remains unknown.

Here, we show that a modest magnetic field (MF) can strongly enhance current generation in living *S. oneidensis* by improving spin-selective electron transfer. The enhancement is large, field-dependent, and occurs without increased biomass or mass transport. Strikingly, the response is asymmetric: it is observed for outward EET to a non-magnetic electrode, disappears for inward electron transfer from the same electrode, but reappears when inward electrons are supplied from a magnetised nickel electrode. These findings indicate that metabolic electrons acquire spin polarisation before entering the EET conduit and are spin-matched in the presence of MF. Consistent with improved electron-transfer efficiency, transcriptomic analysis shows that key EET cytochromes are downregulated under MF exposure despite higher current output, suggesting that fewer electron-transfer components are required when the conduit operates more efficiently.

## EET with magnetic field using non-magnetic electrode

*S. oneidensis* was grown in a three-electrode set-up containing a non-magnetic (carbon felt) working electrode poised at +0.2 V vs. Ag/AgCl (see materials and methods in SM). Under these conditions, the working electrode serves as the terminal electron acceptor, enabling EET from the bacteria to the electrode. The setup was placed between the poles of an electromagnet, with MF perpendicular to the working electrode surface (Fig. 1a). A MF of 0.31 T was applied at two different stages of bacterial growth while performing chronoamperometry (CA). At first, MF was applied after 4 hr of bacterial growth for two 2 hr periods separated by a 1 hr field-free interval. Strikingly, a large MF-induced enhancement in current, with an increase of ∼70 %. In contrast, the corresponding control sample (without MF) showed no appreciable increase in current under the same conditions (Fig. 1b and Fig. S1). This MF-induced response was reproduced in a separate experiment performed at a later growth stage, where field exposure after 26 h of growth enhanced the current by more than 33 % (Fig. 1c and Fig. S2). In both experiments, the current increased gradually throughout the MF exposure period and began to decrease when the MF was switched off between the two intervals. The current enhancement increased linearly with MF strength (Fig. 1d and Fig. S3), indicating a clear field-dependent response and supporting the role of MF exposure in enhancing the current. An abiotic control ruled out any contribution from the growth medium to the MF-induced current response (Fig. S4). Together, these results show that MF exposure of *S. oneidensis* substantially enhances anodic current, which partially decreases upon field removal between exposure intervals and is absent in non-MF-exposed samples.

**Fig. 1.**
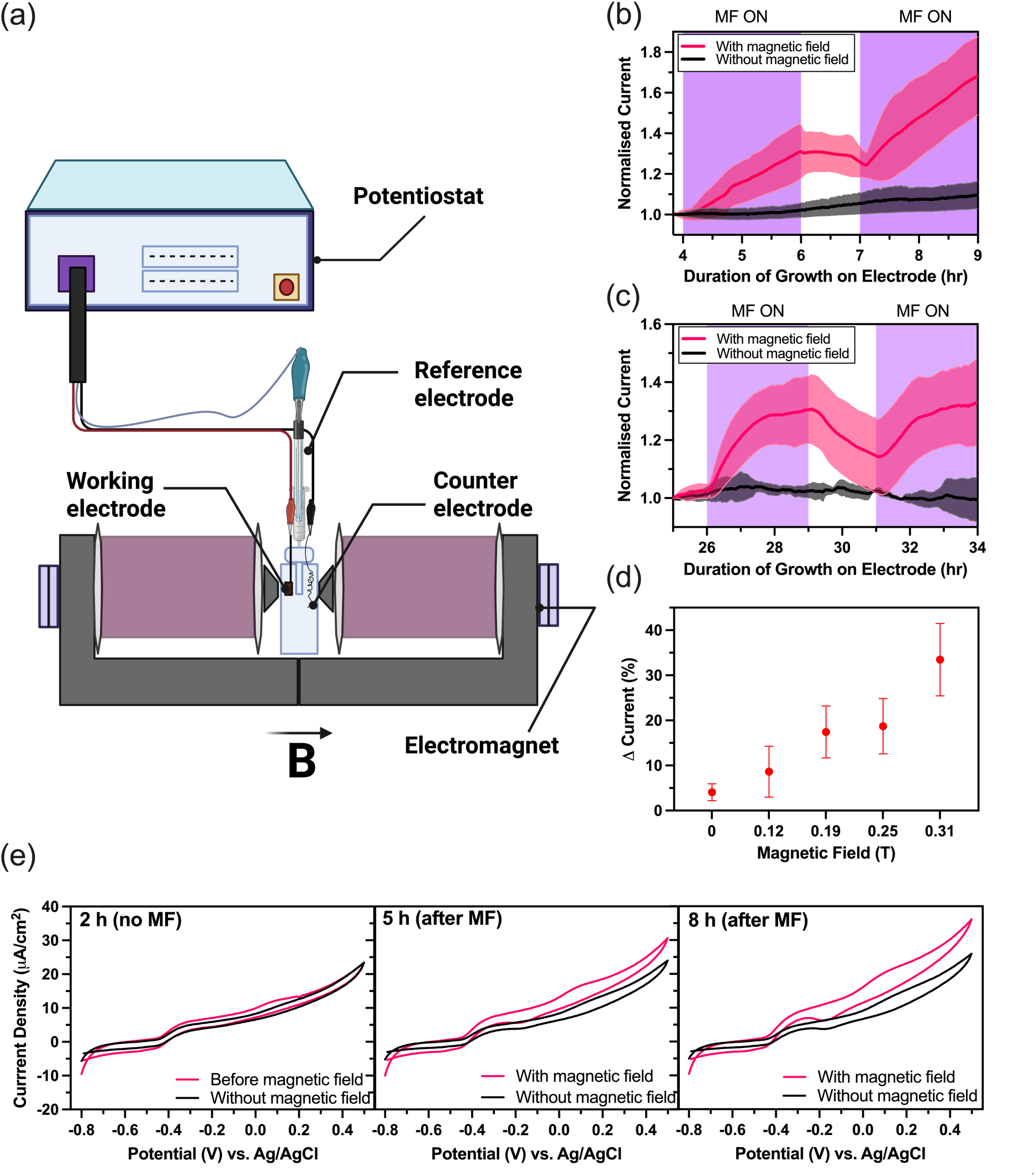
Effect of a magnetic field on extracellular electron transfer to a nonmagnetic electrode. **(a)** Schematic of the experimental setup, consisting of a microbial three-electrode electrochemical cell coupled to an electromagnet. **(b)** Chronoamperometic response of *S. oneidensis* with and without MF exposure using a carbon felt working electrode held at potential of 0.2 V vs. Ag/AgCl (3.5 M KCl). Cells were grown on the electrode for 4 hr before field exposure. Current was normalised to the value recorded 10 min before the first MF exposure. Data show the mean of three biological replicates; error bars, shown as colored shading, represent standard deviation from the mean. **(c)** Chronoamperometric response of *S. oneidensis* under the same conditions after 34 hr of growth on the electrode. Current was normalised to the value recorded 1 hr before the first magnetic-field exposure. Data show the mean of three biological replicates. **(d)** Change in current output vs MF strength applied for 2 hr after 7 hr of growth. Data show the mean of three biological replicates. For (b-d) error bars, shown as colored shading (b, c), represent standard deviation from the mean. The purple shade represents the period when MF was turned on. **(e)** Representative cyclic voltammograms of *S. oneidensis* during MF exposure. Voltammograms were recorded after 2 hr of growth on the electrode (before MF exposure), after 5 hours of growth (after 3 hr of MF exposure) and after 8 hours of growth (after total 6 hr of MF exposure). Scan rate is 5 mV/s.

Cyclic voltammetry (CV) performed on the MF-treated and control samples showed nearly overlapping CV profiles before MF exposure, confirming similar initial electrochemical behaviour that is characteristic of *S. oneidensis* (*26–28*) (Fig. 1e, corresponding CA in Fig. S5). After MF exposure, the CVs diverged, with the MF-treated system showing higher anodic current density, particularly near the cytochrome-based EET region at ∼ +0.2 V (*28*).

Biofilm biomass and planktonic cell density were comparable between MF-treated and control samples as confirmed by crystal violet staining and optical density measurements, respectively (Fig. 2a). Scanning electron microscopy (SEM) after CA showed comparable cell coverage and biofilm morphology on MF-treated and control electrodes, with similar attached cells and nanowire-like structures (Fig. 2b and Fig. S6). Mechanical stirring did not produce a current increase comparable to MF exposure, indicating that magnetohydrodynamic (MHD) effect (*29*, *30*) was not responsible for the MF-dependent response (Fig. S7). Together, these experiments indicate that the MF-induced current enhancement in *S. oneidensis* does not arise from increased biomass or mass-transfer effects, but instead points to improved EET at the biofilm–electrode interface.

**Fig. 2.**
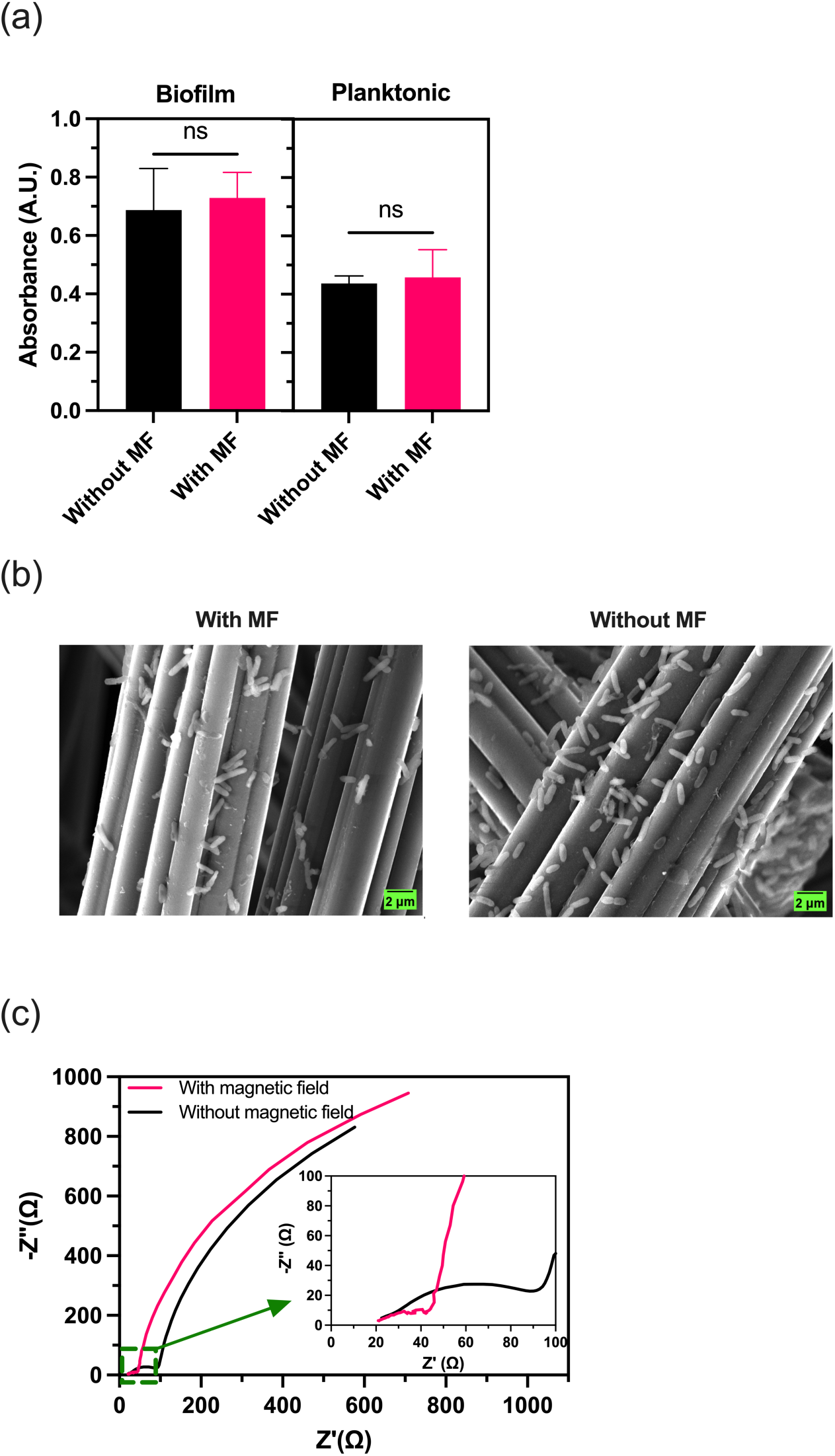
Comparison of cell growth and electrochemical impedance spectroscopy with and without magnetic field. **(a)** Biofilm and planktonic cell quantification with and without MF. Data show the mean of three biological replicates; error bars represent standard deviation from the mean. **(b)** Representative scanning electron micrographs showing comparable *S. oneidensis* cell morphology and electrode colonisation with and without MF. Electrodes are corresponding to the experiment in Fig. 1c. **(c)** Representative Nyquist plot from electrochemical impedance spectroscopy at the potential associated with cytochrome-based electron transfer (200 mV). Inset shows improved electron transfer in the presence of MF.

To validate whether the MF-induced current enhancement was associated with more efficient EET, electrochemical impedance spectroscopy (EIS) was performed. At the potential of the cytochromes (∼ +0.2 V), the Nyquist plots showed smaller semicircles for the MF-treated samples than for the controls, indicating reduced interfacial charge-transfer resistance (Fig. 2c). Equivalent circuit fitting confirmed that the solution resistance remained nearly unchanged, while the charge-transfer resistance at +0.2 V decreased ∼ 2.5-3 times with MF treatment (Supplementary Text-1, and Fig. S8). The results suggest that the dominant effect of MF exposure was on cytochrome-based electron transfer at the biofilm/electrode interface.

We next examined inward EET by performing CA at −0.6 V, where electrons are driven from the non-magnetic electrode toward *S. oneidensis* cells (*31*). In contrast to outward EET, MF exposure produced no measurable change in current (Fig. 3a & Fig. S9a,b, abiotic control in Fig. S9c). Both the control and MF-treated samples showed nearly identical stable current profiles throughout the measurement period. Together, these results reveal an asymmetric response on the non-magnetic electrode: MF enhances outward EET from *S. oneidensis* to the electrode, but does not produce a comparable effect when electron flow is driven inward from the electrode to the cells. This asymmetric response also disfavours any large MHD effect in our system and provides an important mechanistic clue: the MF-induced enhancement is not governed by MF exposure alone, but requires a specific electron-transfer condition which is present only during outward EET.

**Fig. 3.**
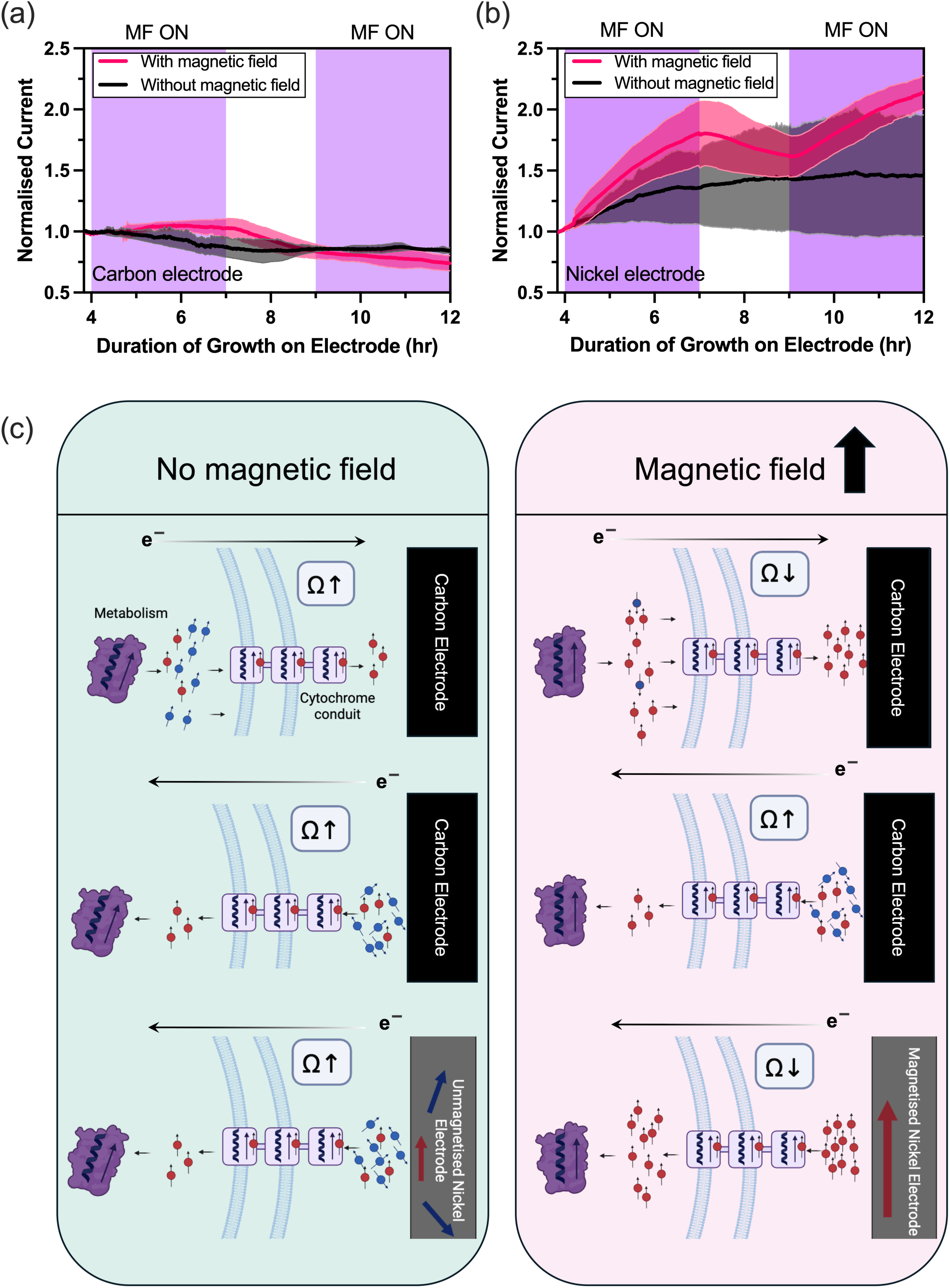
Chronoamperometric analysis of inward EET and schematic model of CISS-enabled MF enhancement. **(a, b)** Chronoamperometric response recording electron uptake by *S. oneidensis* from (a) carbon and (b) nickel electrode poised at −0.6 V with and without magnetic field. The average of three biological replicates is shown in each case. Error bars (coloured shading) refer to standard deviation from the mean. **(c)** Schematic showing electron transfer through the spin-selective cytochrome conduit for carbon and nickel electrodes in the absence and presence of MF (direction indicated by broad black arrow). Red and blue spheres with arrows represent electrons with spin states. With red representing spins states that are matched with the spin-selection axis of the cytochrome conduit. In the absence of MF, the majority of the input spins to the cytochrome conduit are not matched to the spin-selective transport pathway, resulting in greater scattering and higher resistance for the three cases. Under MF exposure, spin matching occurs for outward EET to carbon electrodes and inward electron uptake from magnetised nickel electrodes, leading to reduced resistance and enhanced electron flow.

Recent studies have shown that several cytochromes involved in EET can exhibit spin-selective transport because of their chiral structure (*24*, *25*). This implies that an electron flux that is not spin-matched to the cytochrome’s spin-selective axis will experience higher resistance, whereas a spin-matched flux should pass through the conduit more efficiently (*1–5*, *32*). In *S. oneidensis*, chains of cytochromes form a chiral electron-transfer conduit that can support electron flow in both directions: from intracellular metabolism to the electrode during outward EET and *vice versa* (*16*, *31*). Thus, like the cytochromes that form it, the EET conduit is expected to show lower resistance for a spin-matched electron flux and higher resistance for a spin-unmatched flux.

The applied MF is too weak to directly polarise free electrons. At 0.31 T, the Zeeman splitting is only ∼0.036 meV, far below the room-temperature thermal energy of ∼26 meV, making significant free-electron spin polarisation unlikely. Efficient transport through the spin-selective cytochrome conduit must therefore depend on spin polarisation generated in a prior electron-transfer step, before electrons enter the conduit. Because these upstream steps differ between outward and inward EET, this explains why MF enhances outward electron flow but not inward electron uptake from a non-magnetic electrode.

For inward electron transfer from the non-magnetic electrode to the cells, the incoming electrons are expected to be largely unpolarised, because the electrode itself does not provide spin-selective injection and the applied MF is too weak to directly polarise free electrons. Thus, with or without MF exposure, these unpolarised electrons, being spin mismatched with the conduit, would encounter relatively high resistance, which is what we observe (Fig. 3a).

In contrast, during outward EET, the electrons entering the conduit originate from bacterial metabolism, which involves multiple chiral proteins (*33*, *34*). These upstream metabolic steps may generate a partially spin-polarised electron flux. However, in the absence of MF, this spin-polarisation will not be optimally matched to the spin-selective axis of the cytochrome conduit, leading to greater scattering and higher effective resistance (*1–5*, *32*). Nevertheless, MF exposure provides a common axis that improves the coupling between the upstream metabolic electron-transfer steps and the chiral cytochrome conduit. As new cells attach and the biofilm continues to develop under the MF exposure, newly formed electron-transfer structures may grow with improved coupling between metabolic electron generation and the cytochrome conduit, resulting in spin matching between the two. This would allow electrons with a favourable spin character to pass more efficiently through the conduit, lowering the effective charge-transfer resistance and increasing the anodic current. The gradual rise in current during MF exposure (Fig. 1b,c) further argues against an instantaneous spin reorientation process. Instead, it suggests that the field influences the system progressively as new cells attach and grow. In contrast, when the MF is turned off, newly formed metabolic and cytochrome proteins become spin-decoupled again, leading to a decrease in current.

Proteins can exhibit anisotropic magnetic susceptibility, providing a physical basis for weak MF-induced orientation (*35*). When such anisotropic biological structures form or reorganise in an external MF, magnetic torque can favour orientations with lower magnetic energy (*35*, *36*). For example, Astier et al. showed that lysozyme, bovine pancreatic trypsin inhibitor, and porcine pancreatic α-amylase crystals grown under MF became oriented (*36*). Similarly, Quan et al. reported that bacterial cellulose, a polysaccharide, produced under static MF of 0.045–0.14 T developed a more aligned porous structure (*37*), and Mhamdi et al. showed that a 0.5 T static MF influenced *E. coli* adhesion and orientation on surfaces (*38*). These studies provide precedent that MF exposure can bias the organisation of biological structures during growth or assembly. In our system, MF exposure could therefore influence the organisation of metabolic proteins and the cytochrome conduit in *S. oneidensis*, gradually improving spin coupling between metabolic electron generation and outward EET.

The MF-treated and control samples showed comparable coulombic efficiencies, but substrate consumption was higher under MF exposure (∼17%) than in the control (∼8 %), along with higher current output (Fig. S10). This suggests that MF exposure does not substantially alter the overall electron recovery balance, but increases the rate at which metabolically generated electrons are transferred to the electrode. In other words, the MF appears to relieve a bottleneck in the metabolism–cytochrome–electrode transfer pathway, allowing faster substrate turnover and higher current generation.

## EET with magnetic field using magnetic electrode

To directly test whether a spin-polarised electron current at the input of cytochrome conduit is a key requirement for MF-enhanced EET, we repeated the inward EET experiment using a magnetic nickel foam electrode at −0.6 V. This experiment provided a decisive contrast to the non-magnetic carbon electrode. Nickel is ferromagnetic and becomes magnetised under a modest applied MF, therefore, electrons injected from the nickel electrode under MF are spin polarised (*39*). In the absence of MF, the multidomain magnetic state of nickel would largely average out the spin polarisation of the injected current, making the effective input electron flux comparable to that from a non-magnetic carbon electrode. Strikingly, unlike the carbon electrode, the nickel electrode showed a pronounced increase in cathodic current during inward EET when MF was applied, with current enhancements of > 100 % (Fig 3b & Fig.S11a-c; abiotic control in Fig. S11d). This result strongly supports our hypothesis that MF enhances EET by spin-matching input electron flux to the chiral cytochrome conduit.

The proposed mechanism for all the three cases (with and without MF) is illustrated in Fig. 3c. It has a conceptual analogy with spintronic devices such as spin valves and magnetic tunnel junctions, where charge transport is not governed only by charge flow, but also by the spin state of the electrons (*32*, *40–42*). In these devices, one magnetic layer acts as a spin polariser, creating a spin-biased current, while another layer acts as a spin analyser or detector. The resistance then depends on how well the spin orientation of the transmitted electrons matches the magnetic configuration of the receiving layer. When the spin polarisation is favourably aligned, electrons are transmitted more efficiently and the resistance is lower; when the spin orientation is mismatched, scattering increases and the resistance rises.

It should be noted that MF has previously been used as an external stimulus in MFCs, with reported gains in output from *Shewanella-*based and *Geobacter*-based systems, as well as mixed-culture systems (*43–47*), However, these studies interpreted the response mainly through changes in microbial activity and gene expression, biofilm development, internal resistance, electrode effects, or community composition, rather than through spin-dependent electron transport. Thus, they did not establish whether the MF response of live electroactive bacteria originates from spin-selective transport through the chiral EET conduit, which is the central question addressed in our work.

The non-magnetic carbon felt electrode is porous, allowing bacterial attachment and biofilm growth on fibres with different spatial orientations. To examine MF-direction effects more directly, bacteria were grown on one side of a graphite electrode (Fig. 3c), and the MF was applied in either the positive or negative direction. Anodic current increased in both cases, although the enhancement under the positive MF direction was slightly higher, by approximately 4% (Fig. 4). This weak polarity-dependent response supports the link between MF-induced current enhancement and the CISS character of chiral proteins in the EET pathway (*24*, *25*). However, because bacterial cells are heterogeneously oriented at the electrode surface, the present setup does not allow the exact field-orientation dependence of EET current to be precisely measured.

**Fig. 4.**
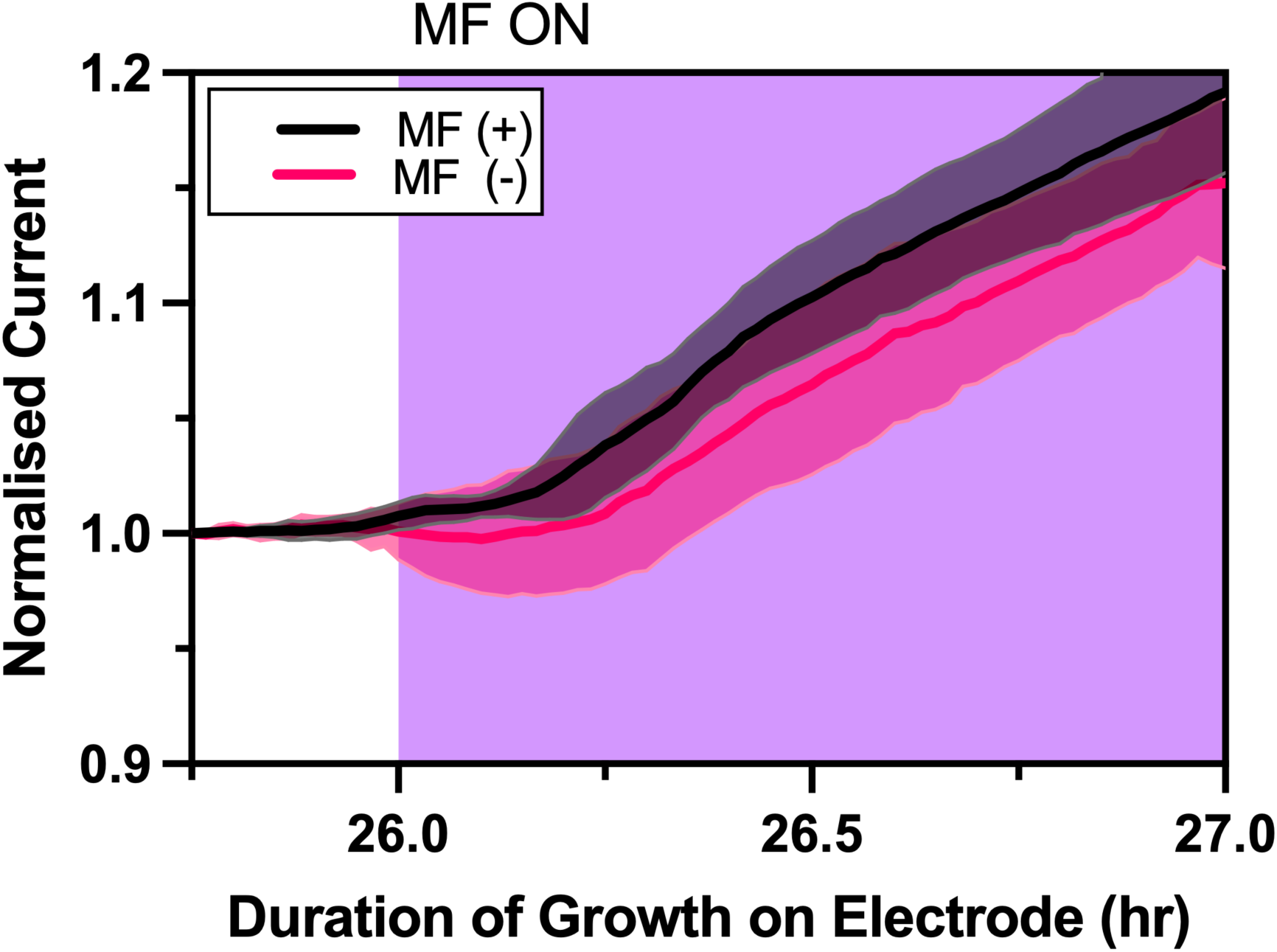
Effect of magnetic field orientation. Chronoamperometric response at +0.2 V for different orientation of magnetic field. The average of five biological replicates is shown in each case. Error bars (coloured shading) refer to standard deviation from the mean.

## Differential gene expression in magnetic field

RNA-seq analysis of MF-treated and control samples revealed a clear transcriptional response to MF exposure. Principal component analysis (PCA) separated the two conditions along PC1, and volcano plot identified distinct sets of significantly upregulated and downregulated genes across a broad range of transcript abundances, as indicated by MA plot (Fig. 5a,b and Fig. S12). Hierarchical clustering of the most differentially expressed genes further confirmed treatment-specific grouping (Fig. S13). GO and KEGG enrichment analyses showed that MF exposure produces broad transcriptional remodeling in *S. oneidensis*, affecting nucleic-acid-related processes, protein export, quorum sensing, motility, and central metabolism (Fig. S14 and Fig. S15).

**Fig. 5.**
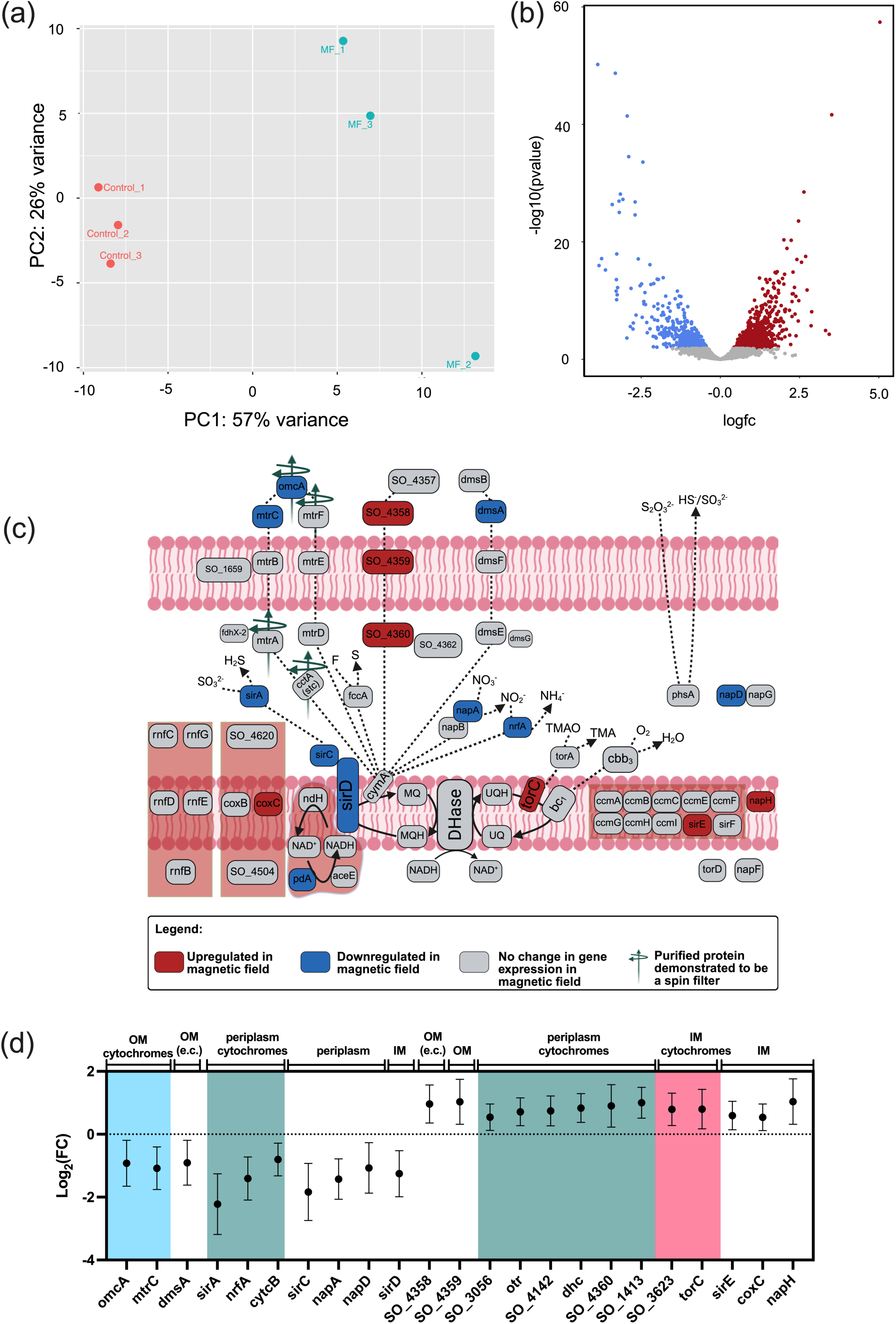
Differential gene expression in presence of magnetic field during extracellular electron transfer. **(a)** PCA plot showing distinct differences in MF and control samples. **(b)** Volcano plot showing differential gene expression across conditions. **(c)** Differentially expressed EET-related genes, with red indicating upregulation, blue indicating downregulation and grey indicating no change in expression in MF. Proteins that have previously been demonstrated to be spin selective are indicated by green arrows. Details of genes, including references for cellular location, are included in Table S2. **(d)** Log_2_(FC) of EET genes in MF. Error bars represent 95% confidence intervals calculated from the standard error. Blue panel indicates outer-membrane cytochromes, green panel indicates periplasmic cytochromes and pink panel indicates inner-membrane cytochromes. OM (e.c.) = outer membrane (extracellular).

Gene set enrichment analysis of 481 proteins (*48*) implicated in EET in *S. oneidensis* shows there is overall a significant downregulation of electron transfer proteins (Fig. S16), yet there is enhanced current output. A visualization of the differentially expressed genes is shown in Fig. 5c, highlighting that differential expression effects all levels of the EET pathway: from the inner membrane, across the periplasm, to the outer membrane. Of the 42 cytochromes encoded in *S. oneidensis* (*49*), 13 are differentially expressed in MF. While certain periplasmic and inner-membrane cytochromes were upregulated, remarkably, the major outer-membrane cytochromes *mtrC* and *omcA* were downregulated in MF (Fig. 5d and Table S2). There were no other outer-membrane cytochromes upregulated to compensate for this. It should be noted that OmcA has already been shown to be spin selective (*25*). Although the less well-characterised SO_4358–SO_4360 locus was upregulated under MF exposure, annotation shows SO_4358 is not a cytochrome, despite being comparable in size to MtrC and MtrF (*50*), and therefore cannot compensate for the reduced expression of outer-membrane cytochromes. Despite downregulation, all the above genes are still expressed as seen from the normalised counts, just at different levels in the MF (Fig. S17).

The transcriptomic result is counterintuitive: MF exposure increases current output even though several key EET components are downregulated. Since no compensatory upregulation of alternative outer-membrane cytochromes was observed, the enhanced current cannot be explained by increased production of such EET proteins. Instead, these data suggest that MF exposure improves the efficiency of the existing EET pathway, allowing a reduced cytochrome network to transfer electrons more effectively. In other words, under MF exposure, each active EET conduit may carry a higher electron flux, thereby reducing the need for higher expression of EET components in *S. oneidensis*.

Together, our results identify spin selectivity as a key determinant of EET efficiency in live *S. oneidensis*. MF exposure enhances electron flow through the chiral cytochrome conduit by spin-matching. This is revealed by the directional response: outward metabolic electron flow is enhanced on non-magnetic electrodes, inward electron uptake from the same electrode is not, and the enhancement is restored when electrons are injected from magnetised nickel. This asymmetry also implies that metabolic electrons are already spin-polarised before they enter the EET conduit. More broadly, these findings extend CISS from purified biomolecules to a functional process in living cells and suggest that, because proteins are intrinsically chiral, spin selectivity may be an overlooked feature of biological electron transport beyond electroactive bacteria. Practically, MF can be used to modulate microbial current generation, providing a spin-based strategy to improve charge extraction in bioelectrochemical systems.

## Acknowledgements

The work is supported by Anusandhan National Research Foundation, India (Grant No. CRG/2023/002839). The authors thank Central Research Facility (CRF), IIT Delhi, for use of the SEM facility.

## Author contributions

Conceptualization: LED, RM

Methodology: LED, RM, KK

Investigation: KK

Visualization: LED, RM, KK

Funding acquisition: LED, RM

Project administration: LED, RM

Supervision: LED, RM

Writing – original draft: KK, RM, LED

Writing – review & editing: LED, RM

