## Supplementary Materials for "Magnetic field enables spin matching in spin-selective pathways to enhance current in bacteria"

**for**

### Materials and methods

#### Growth of *Shewanella oneidensis*

*Shewanella oneidensis* MR-1 cells (ATCC) were grown in LB broth under aerobic conditions and 150 rpm shaking at 30 °C. The culture was then adjusted to optical density (at 600 nm) of 1.0 with sterile modified M1 growth medium to maintain consistent cell numbers. Three mL of this adjusted culture was then added to 12 mL of the same growth medium, for a final volume of 15 ml in the sterile electrochemical cells.

Electrochemical experiments used the following growth medium which consisted of a base medium (*1*) and trace vitamins, minerals and amino acids (*2*). (The base medium consisted of the following (per litre): HEPES, 7.2 g; NaOH, 0.3g; $NH_4Cl$, 1.5 g; KCl, 0.1 g; $NaH_2PO_4$, 0.52 g, followed by filter-sterilised minerals, vitamins, amino acids and $CaCl_2$, 7.5g. The trace mineral stock consists of the following (per litre): $C_6H_9NO_6$, 1.5 g; $MgSO_4.H_2O$, 3 g; $MnSO_4.H_2O$, 0.5 g; NaCl, 0.5 g; $FeSO_4.7H_2O$, 0.5 g; $CaCl_2.2H_2O$, 0.1 g; $CoCl_2.5H_2O$, 0.1 g; $ZnCl_2$, 0.13 g; $CuSO_4.5H_2O$, 0.01 g; $AlK(SO_4)_2.12H_2O$, 0.01 g; $H_3BO_3$, 0.01 g; $Na_2MoO_4.2H_2O$, 0.025 g; $NiCl_2.6H_2O$, 0.024 g, $Na_2WO_4.2H_2O$, 0.025 g. The vitamin stock consists of the following (per litre): biotin (D-biotin), 2 mg; olic acid, 2 mg; pyridoxine HCl, 10 mg; riboflavin, 2 mg; thiamine HCl 0.1 H2O, 5 mg; nicotinic acid, 5 mg; D-pantothenic acid, hemicalcium salt, 5mg; B12, 0.1mg; p-aminobenzoic acid, 5mg; thioctic acid, 5mg. The amino acid stock consists of the following composition (per litre): L-glutamic acid, 2 g; L-arginine, 2 g; DL-serine, 2 g.

For all experiments, 20 mM sodium lactate was added to the above as carbon source. For experiments analysing inward EET from an electrode (i.e. electron consumption by *S. oneidensis*), the same medium was used to support growth, with the additional step of $N_2$ sparging and 50 mM fumarate added as the terminal electron acceptor (*3*).

#### Electrochemical set-up

Depending on the experiment, the working electrode consisted of either carbon felt, nickel foam or graphite (indicated in each case). For all experiments, a titanium wire (Sigma Aldrich) was used as a counter electrode, and an Ag/AgCl (3.5 M KCl) was used as the reference electrode. All potentials in this work are reported with respect to this reference electrode. A frit was attached to a capillary with the help of heat shrink, and a salt bridge was used to connect the reference electrode to the electrochemical cell. The set-up connected to a potentiostat (MultiPalmSens 4, Palmsens) to record the data. Where indicated, a magnetic field (MF) was provided with an electromagnet (Model EM-17, Raman Scientific Instruments) The MF was calibrated using a gaussmeter. Unless otherwise specified, the MF applied was 0.31 T.

#### Chronoamperometry using various electrodes

For experiments using a non-magnetic electrode, carbon felt (Sigma Aldrich) that was used as a working electrode with dimensions 1 cm x 1 cm x 0.3 cm (geometric surface area = 3.2 $cm^2$). Prior to use, these electrodes had been stored in 1 M HCl and were subsequently washed with distilled water and autoclaved before use. For experiments measuring outward EET experiments, the carbon felt was poised at +0.2 V during chronoamperometry (CA). For inward EET experiments, it was poised at −0.6 V during CA.

For experiments using a magnetic electrode, nickel foam with dimensions 1 cm x 1 cm x 0.1 cm (geometric surface area = 2.4 $cm^2$) was used as the working electrode. The electrodes were prepared using a modified version of a previously reported protocol to remove the oxide layer (*4*). Briefly, thee electrodes were dipped in acetone for 20 minutes and transferred to 1 M HCl for 10 minutes. The electrodes were subsequently washed with distilled water and autoclaved before use. The nickel electrodes were poised at −0.6 V during CA.

For experiments investigating MF orientation, graphite electrodes of the dimensions 1 cm x 1 cm x 0.2 cm (geometric surface area = 2.8 $cm^2$) were first polished with sandpaper of grit P1000 to make the surface uniform and rough enough so that bacteria can adhere to it. Then the electrodes were kept in 1 M HCl overnight subsequently washed with distilled water and autoclaved before use. Further, to keep only one side exposed to the bacterial attachment, autoclave tape covered all 5 remaining sides of the electrode.

All CA was performed on biological triplicates at minimum.

**Cyclic voltammetry and electrochemical impedance spectroscopy**
Cyclic voltammetry (CV) was performed in a potential scan window from −0.8 V to +0.5V with scan rate 5 mV/s.

The following parameters were using for electrochemical impedance spectroscopy (EIS) on samples that were grown for 26 hr on the carbon felt electrode, followed by 2 hr MF: T (equilibration) = 10 s, E (initial) = –0.8 V, E (final) = +0.5 V, decade = 12, E(step) = 0.01V, E(ac) (sinus amplitude) = 0.05 V and frequency range: 1 to $10^5$ Hz. The two potentials extracted were −0.4V and +0.2V, as they depict the flavin and cytochrome behaviour, respectively,. The Nyquist plots were then analysed and fitted into the circuit to generate parameters using MultiTrace 4.5.

All CV and EIS were performed on biological triplicates at minimum.

**Scanning electron microscopy**
Electrodes in MF treated and control experiments were stored overnight in 2.5% glutaraldehyde and subsequently treated as reported in (*5*). Briefly, this consisted of serial dehydration with acetone (with 25, 50, 75, and 100% concentration) diluted with water, and hexamethyldisilazane (HMDS) (with 25, 50, 75, and 100% concentration) diluted with acetone Each electrode was immersed in acetone and HMDS for 30 minutes, starting from the lowest to the highest concentration. The last step in HMDS was repeated three times to make sure that the samples were appropriately fixed. Finally, the samples were air-dried at room temperature and stored. The SEM images were captured by FESEM TESCAN and JEOL JSM-7800F Prime instruments.

**Quantification of biofilm, planktonic cell growth and lactate consumption**
To quantify biomass on the working electrode, a modified version of the protocol reported in (*6*) was used. Briefly, electrodes were rinsed with sterile distilled water and incubated with 3 mL of 0.1 % crystal violet for 30 minutes. The electrodes were taken out and dipped in 3 mL of distilled water for 10 seconds, followed by destaining with 30 % acetic acid. The optical density of this mixture with 5X dilution with acetic acid were measured at 550 nm to quantify biofilm.

To quantify planktonic cell growth, the optical density of 1 mL of the microbial culture was measured at 600 nm.

Lactate was estimated using Megazyme Lactic Acid Rapid Assay Kit.

**Transcriptomic sample preparation, RNA extraction, library preparation, and sequencing**

*S. oneidensis* was grown in three-electrode set-ups containing carbon felt working electrodes poised at 0.2 V versus Ag/AgCl. Three control and three MF-treated reactors were operated for 10 hr 15 min. The MF was applied in two intervals: from 4 hr to 7 hr and from 7 hr 15 min to 10 hr 15 min, with a 15 min break between exposures.

At the end of the experiment, 10 mL PBS was added to each setup and the biomass was harvested by centrifugation at 4500 rpm for 10 min. The supernatant was discarded, and the tubes were briefly inverted on tissue paper for 10–20 s to remove residual liquid. The pellet was immediately resuspended in 5 mL RNAprotect Bacteria Reagent (Qiagen) by gentle shaking. Samples were centrifuged again at 4500 rpm for 10 min, after which the supernatant was discarded and the tubes were briefly inverted on tissue paper. The final pellets were sealed with parafilm and stored at −80 °C until RNA extraction.

Total RNA was extracted using TRIzol. RNA concentration and integrity were assessed using Qubit and Agilent TapeStation. For rRNA-depleted total RNA sequencing, bacterial RNA was treated with the QIAseq FastSelect −5S/16S/23S Kit (Qiagen) following the manufacturer's instructions. The resulting rRNA-depleted RNA was used for library preparation with the NEBNext Ultra II RNA Library Prep Kit for Illumina (New England Biolabs). Final libraries were assessed using TapeStation and Qubit 4 fluorometer; all libraries passed QC, with average library sizes ranging from 315 to 361 bp. Sequencing was performed on the Illumina NovaSeq 6000 platform to generate approximately 20 million 150 bp paired-end reads per sample.

**RNA-seq data processing and differential expression analysis**

Raw paired-end RNA-seq reads from control and MF-treated samples, with three biological replicates for each condition, were analysed on the Galaxy platform (*7*) using the public usegalaxy.eu server. Initial read quality was assessed using FastQC v0.12.1, and quality-control reports were summarised using MultiQC v1.27 (*8*). Reads were pre-processed using fastp v1.0.1 (*9*) with default quality and length filtering enabled. Bases with Phred quality scores below Q15 were treated as unqualified, and reads were filtered using the default thresholds: maximum 40% unqualified bases per read, maximum 5 ambiguous bases, and minimum read length of 15 nt. Filtered reads were aligned to the *S. oneidensis* MR-1 reference genome, NCBI RefSeq assembly GCF_000146165.2/ASM14616v2, using Bowtie2 v2.5.3 (*10*). Gene-level read counts were generated from the resulting alignments using featureCounts v2.1.1 (*11*) with the corresponding RefSeq genome annotation. Differential gene expression between control and MF-treated samples was analysed using DESeq2 v1.40.2 (*12*). Principal component analysis were performed using DESeq2-normalised counts. Heatmaps were generated using heatmap.2 from variance-stabilising transformation (VST)-normalised counts for genes with $|\log2 \text{ fold change}| > 1$ and Benjamini–Hochberg FDR-adjusted P value $\leq 0.05$, with z-scores computed row-wise prior to clustering. Volcano plots were generated using ggplot2. MA-plot was generated by DESeq2.

**Functional Enrichment and Gene Set Analysis**

Gene Ontology (GO) enrichment analysis was performed using ShinyGO v0.85.1 (*13*) with *Shewanella oneidensis* selected as the reference species using the STRINGdb annotation. Enriched terms were identified using an FDR cutoff of 0.05, with pathway sizes restricted to

2–5000 genes and redundancy removal enabled. For visualisation, the top 20 enriched terms were displayed separately for genes upregulated and downregulated under MF treatment.

KEGG pathway enrichment analysis was performed with KOBAS-I (KOBAS 3.0) (*14*) using default settings. Upregulated and downregulated genes under MF treatment were analysed separately.

Gene Set Enrichment Analysis (GSEA) was performed using the software GSEA 4.4.0 (*15*, *16*) using a custom gene set comprising 481 electron-transfer-associated genes (*17*). GSEA was run using DESeq2-normalised expression counts with the MF and control samples as phenotype classes. Genes were ranked using the signal-to-noise metric, and enrichment was tested using the weighted scoring scheme with 10,000 gene-set permutations.

**Supplementary Text – 1**

EIS spectra from MF-treated and control samples were fitted using the equivalent-circuit model shown in Fig. S8a (*18*, *19*), and representative fits are shown in Fig. S8b-e. The circuit consists of the solution resistance ($R_s$), a Randles element containing the charge-transfer resistance ($R_{ct}$), double-layer capacitance ($C_{dl}$), and Warburg diffusion element (W), together with an additional biofilm interface represented by ($R_B$) and ($C_B$). The mean of fitted values from biological triplicates are summarised in Table S1 for measurements performed at −0.4 V and +0.2 V.

The solution resistance, $R_s$, was consistent across all measurements, regardless of potential, indicating a constant resistance from the M1 growth medium. Symbolically, $R_B$ represents the resistance to electron hopping across the biofilm via electron shuttles or cytochromes, and $C_B$ represents the total localised charge separation of the *Shewanella* biofilm layer. Their values are large for both MF treated and control, indicating biofilm development over the electrodes along with a potentially complex biofilm matrix (*20*).

Interestingly, the resistance to charge transfer, $R_{ct,}$ at −0.4 V and 0.2 V, was two to three times lower for MF-treated samples than the values for the control, establishing that the MF treatment made a significant difference via direct electron transfer through cytochrome and mediated electron transfer via flavins. In addition to acting as electron shuttles (*21*), flavins are involved in electron transfer between MtrC/OmcA and electrodes by binding to these structures (*22*, *23*). Double-layer capacitance, $C_{dl,}$ was lower in MF samples. The Warburg (W) element was included to represent the diffusion phenomenon, but due to an additional biofilm interface, the characteristic Warburg line of 45° was not observed.

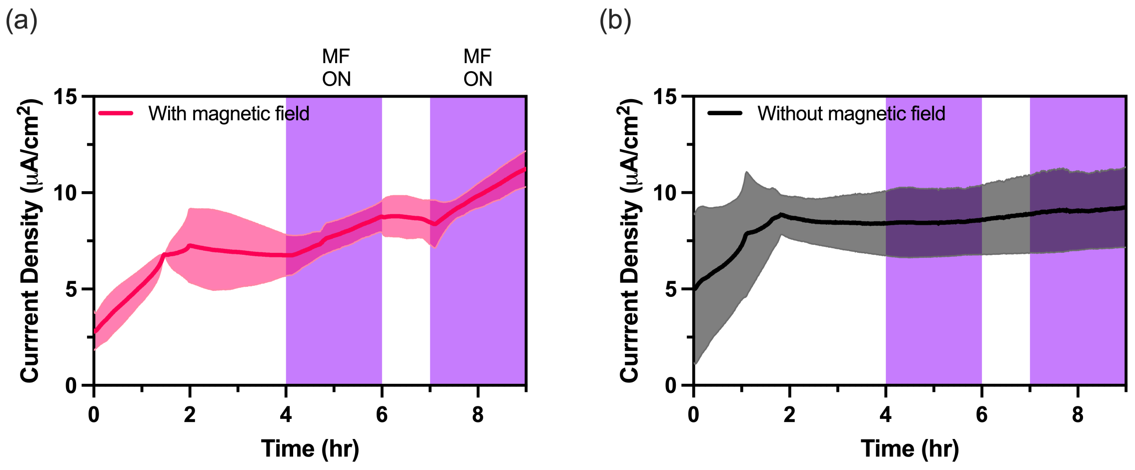


**Fig. S1. (a, b)** CA response of *S. oneidensis* grown on carbon felt poised at +0.2 V (a) with MF and (b) without MF. Data show the mean of three biological replicates; error bars, shown as colored shading, represent standard deviation from the mean. The purple shade represents the period when MF was turned on.

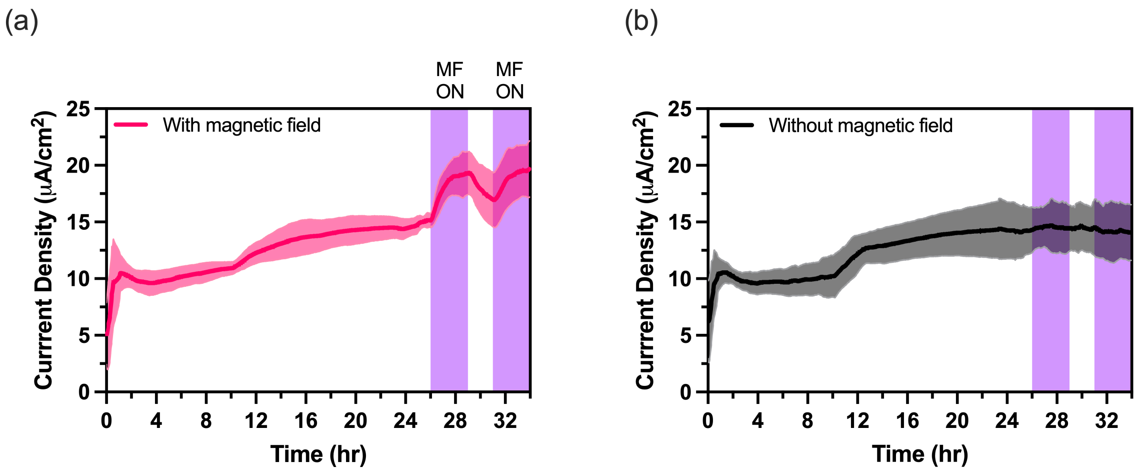


**Fig. S2. (a, b)** CA response of *S. oneidensis* grown on carbon felt poised at +0.2 V (a) with MF and (b) without MF. Data show the mean of three biological replicates; error bars, shown as colored shading, represent standard deviation from the mean. The purple shade represents the period when MF was turned on.

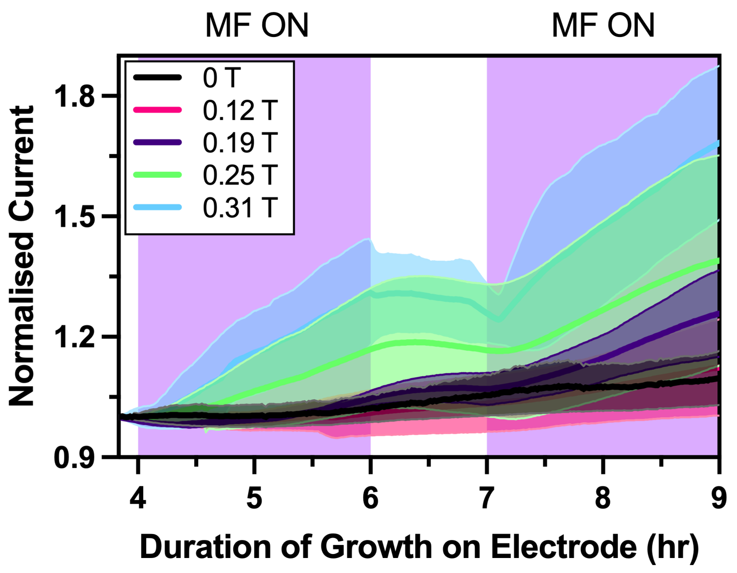


**Fig. S3.** Normalised CA response of *S. oneidensis* grown on carbon felt poised at +0.2 V with various strengths of MF. Data show the mean of three biological replicates; error bars, shown as colored shading, represent standard deviation from the mean. The purple shade represents the period when MF was turned on. Current is normalised to the value 10 min before initial MF exposure.

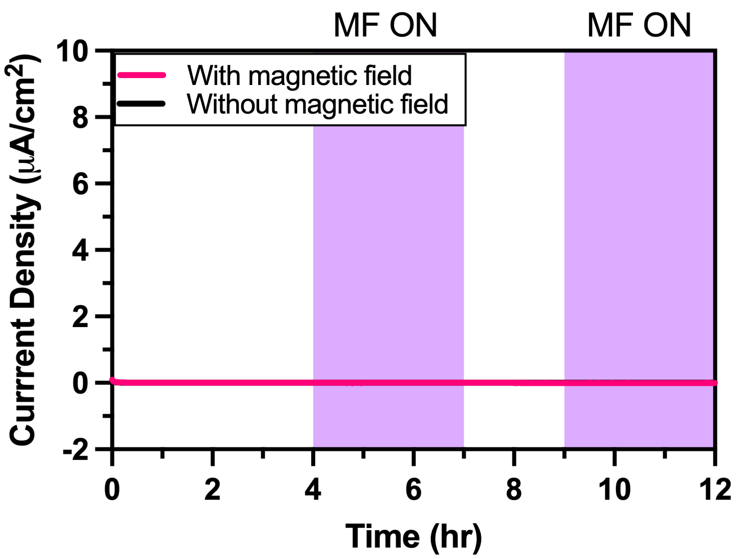


**Fig. S4.** Representative CA response of abiotic control (sterile growth medium) with and without MF using carbon felt poised at +0.2 V. The purple shade represents the period when MF was turned on.

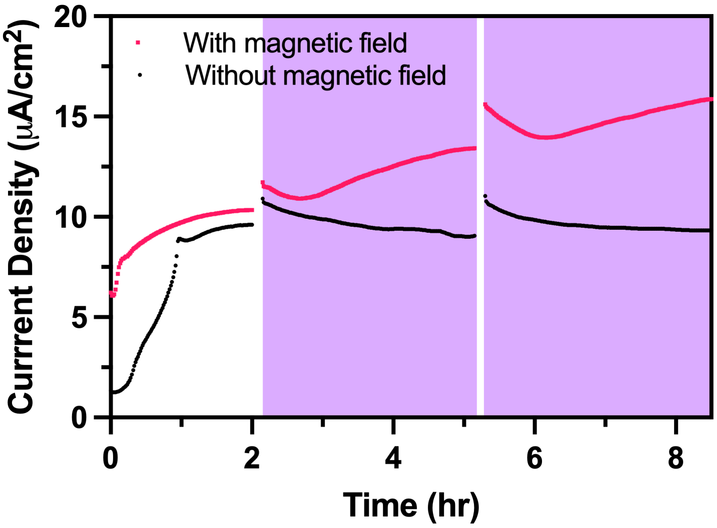


**Fig. S5.** CA response of *S. oneidensis* grown on carbon felt poised at +0.2 V with and without MF exposure during the CV measurement. Breaks in the graph represent the period when CV was conducted. The purple shade represents the period when MF was turned on.

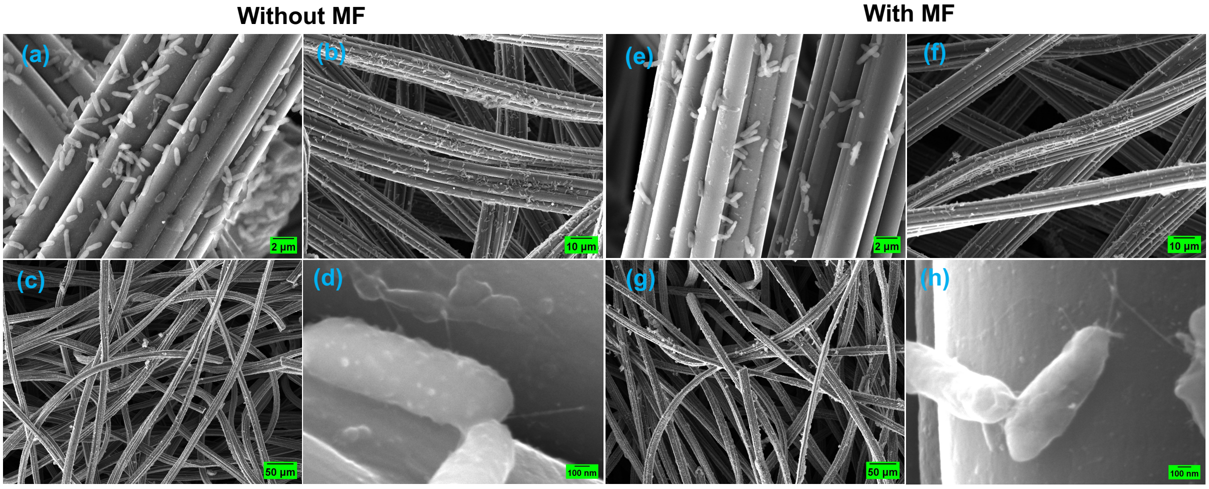


**Fig. S6.** Representative scanning electron micrographs at various magnifications of *S. oneidensis* with and without MF on the carbon felt electrodes after the measurements in Fig. S2.

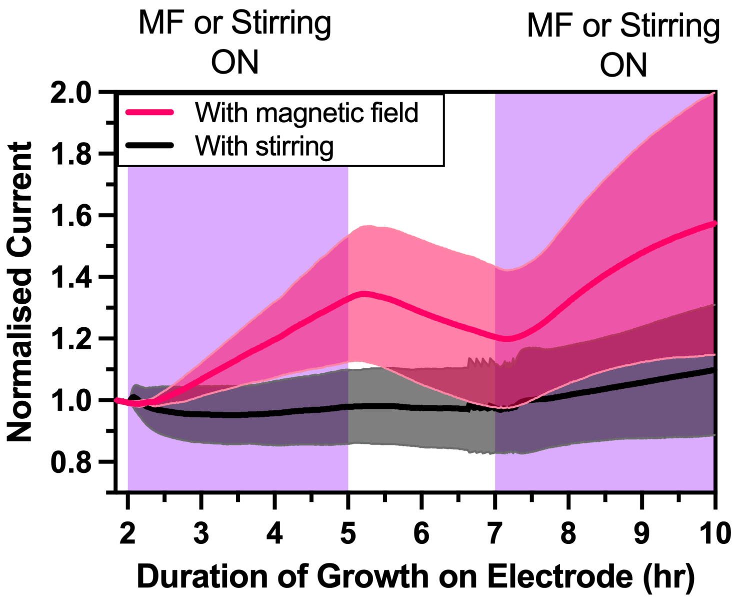


**Fig. S7.** Normalised CA response of *S. oneidensis* grown on carbon felt poised at +0.2 V with MF and with stirring. Data show the mean of three biological replicates; error bars, shown as colored shading, represent standard deviation from the mean. The purple shade represents the period when either MF or stirring was turned on. Current is normalised to the value 10 min before initial MF exposure or stirring.

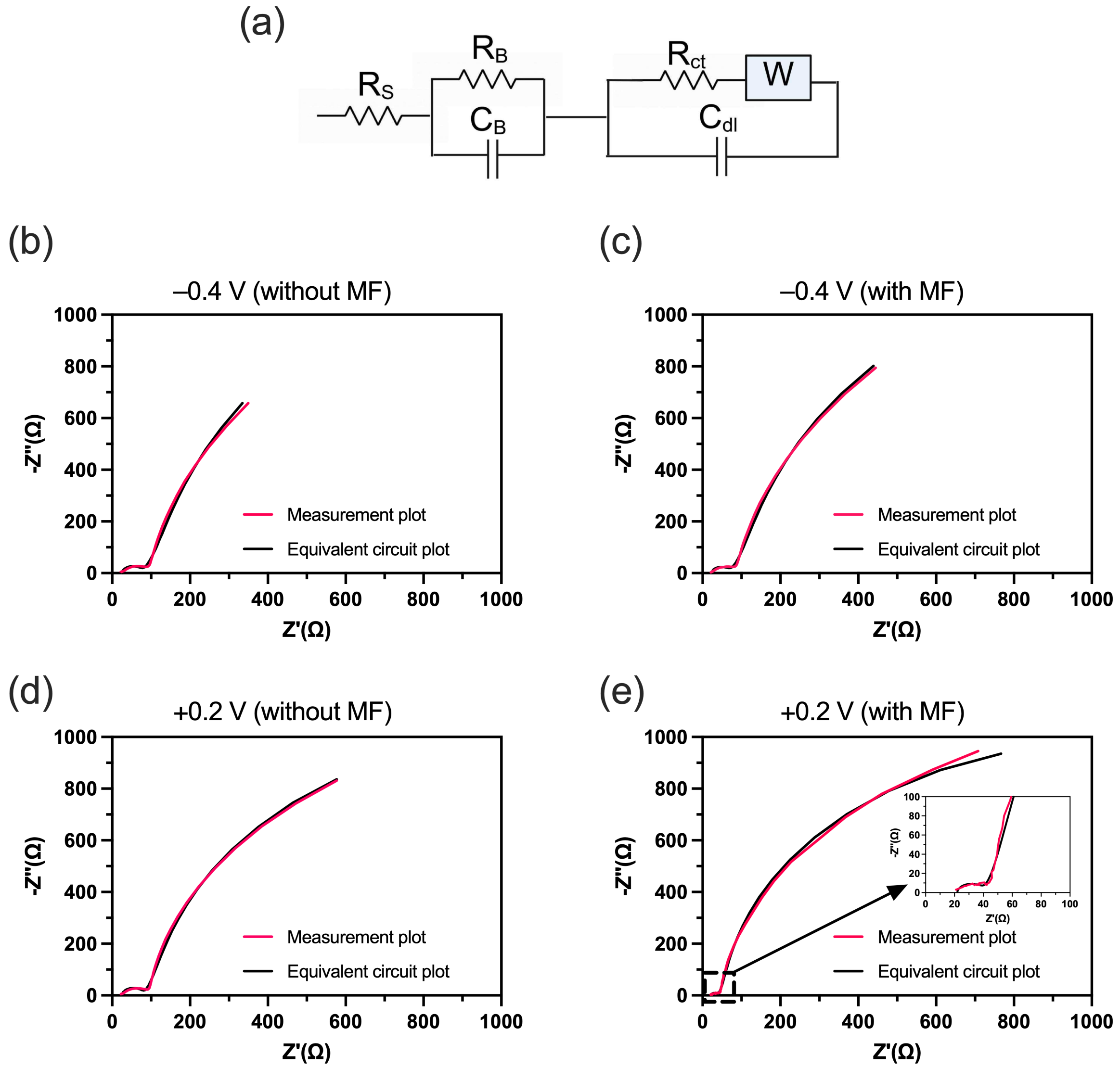


**Fig. S8.** (a) Equivalent circuit model used for fitting the EIS spectra. (b-e) Circuit fitted plots of EIS of *S. oneidensis* grown on carbon felt poised at +0.2 V and −0.4V with and without MF. Cells were grown for 26 hr on the electrode, followed by 2 hr of MF before performing EIS.

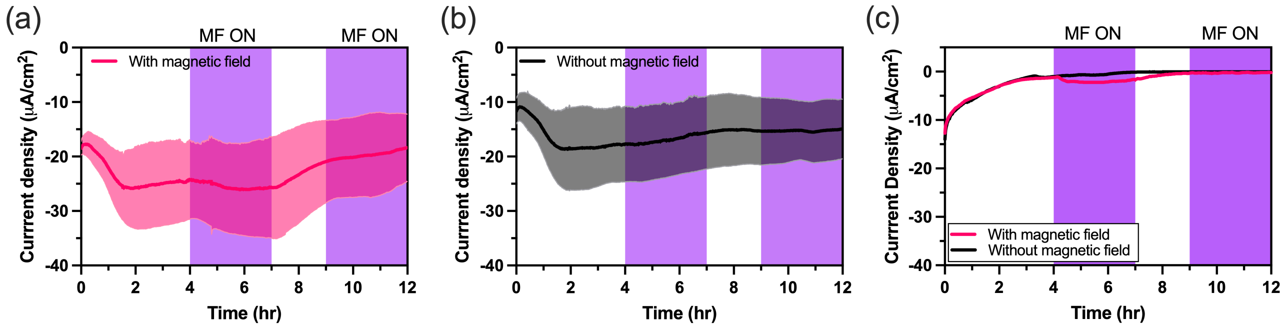


**Fig. S9.** (a,b) CA response of *S. oneidensis* grown on carbon felt poised at −0.6 V (a) with and (b) without MF. Data show the mean of three biological replicates; error bars, shown as colored shading, represent standard deviation from the mean. The purple shade represents the period when MF was turned on. (c) Representative abiotic control (sterile growth medium) under the same condition with insignificant current recorded.

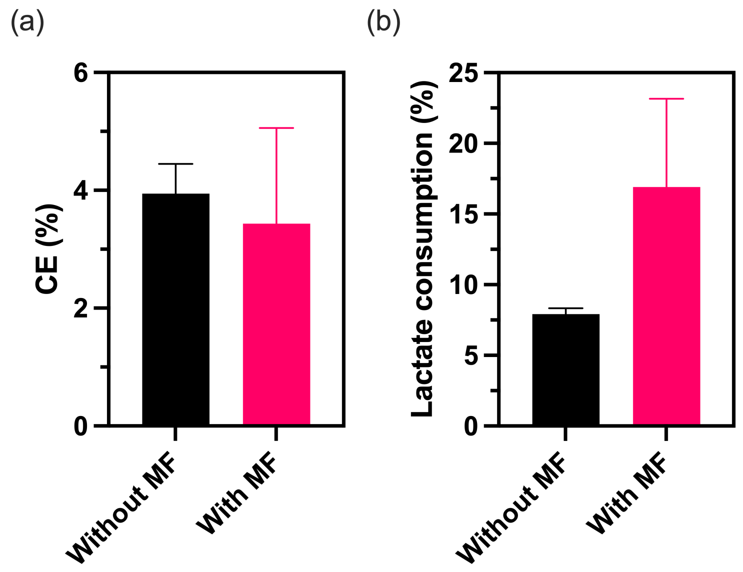


**Fig. S10.** (a) Coulombic efficiency (CE) with and without MF, assuming lactate is degraded to acetate. (b) Fraction of lactate consumed with and without MF. *S. oneidensis* was grown on carbon felt poised at +0.2 V for 4 hr prior to a total MF exposure of 6 hr.

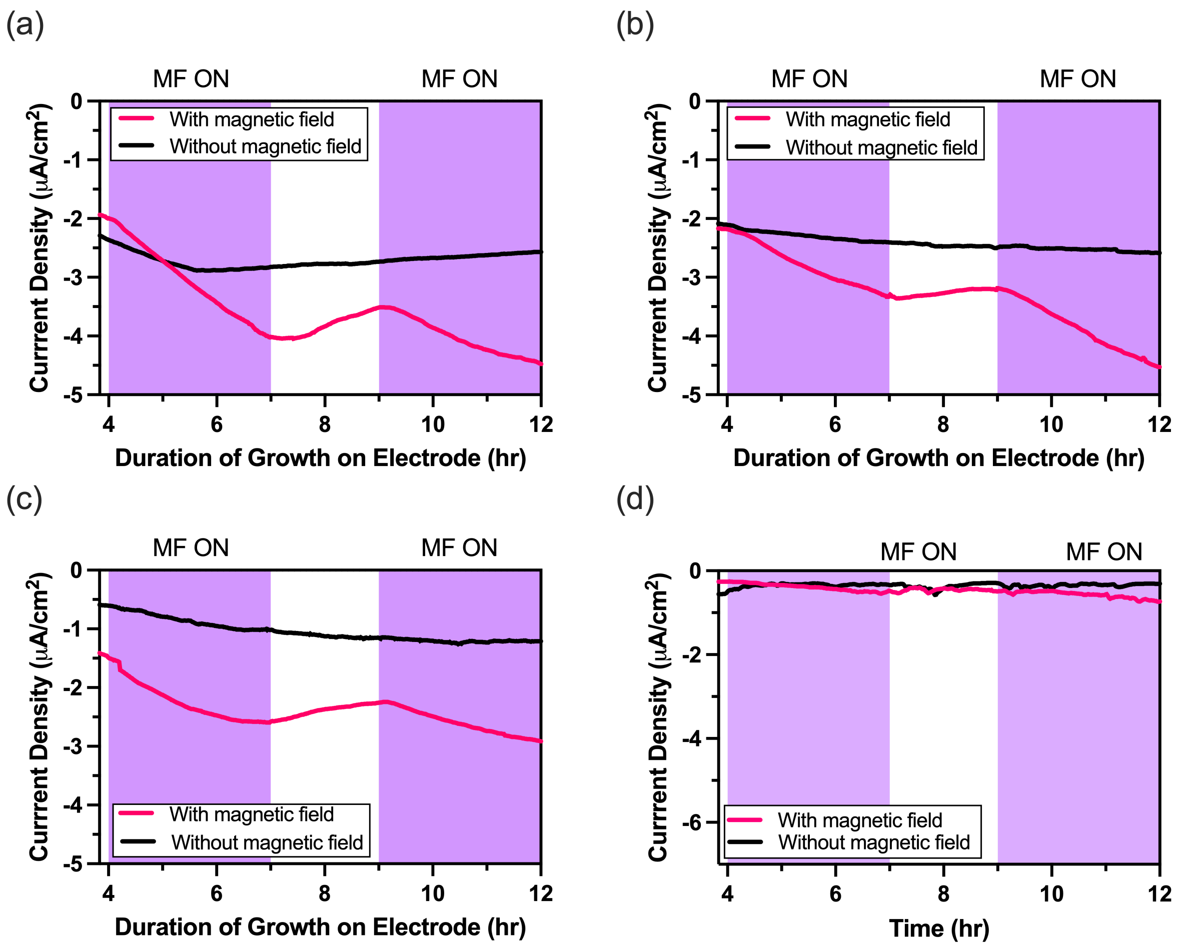


**Fig. S11.** (a-c) Individual biological replicates of *S. oneidensis* grown on nickel electrode at –0.6 V with and without MF. The purple shade represents the period when MF was turned on. (d) Representative abiotic control (sterile growth medium) under the same condition with insignificant current recorded.

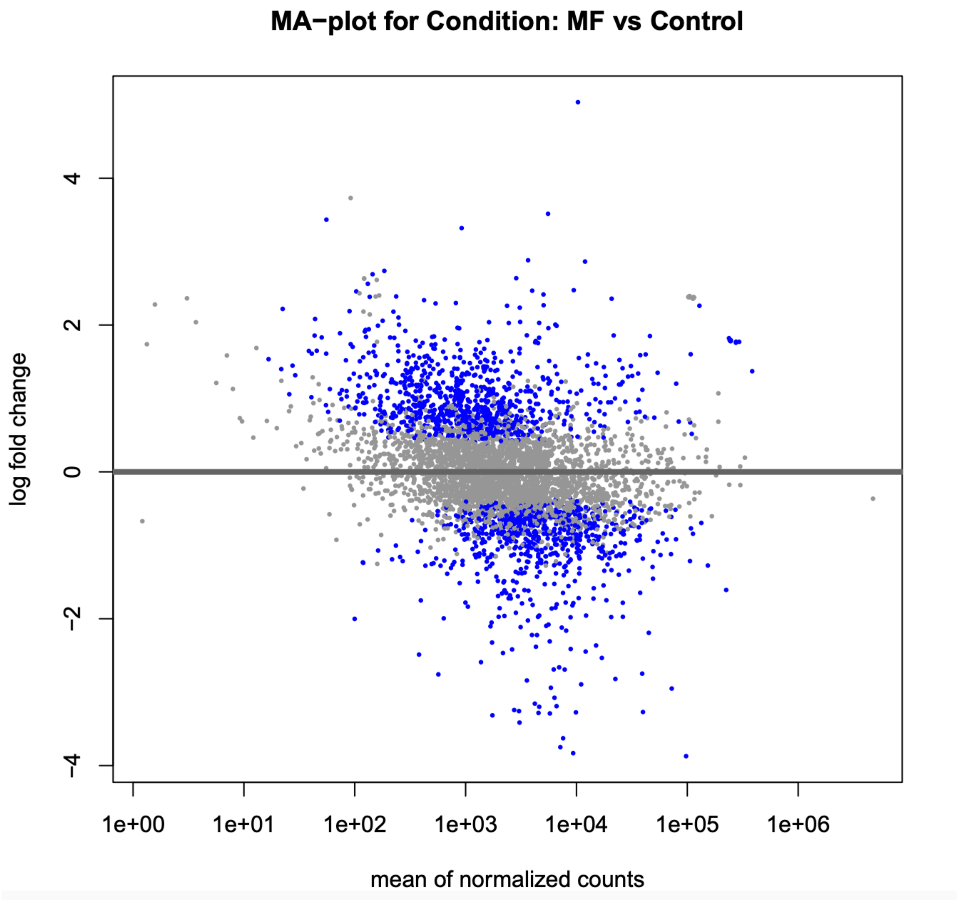


**Fig. S12.** MA plot showing differentially expressed genes are distributed across a broad range of transcript abundances.

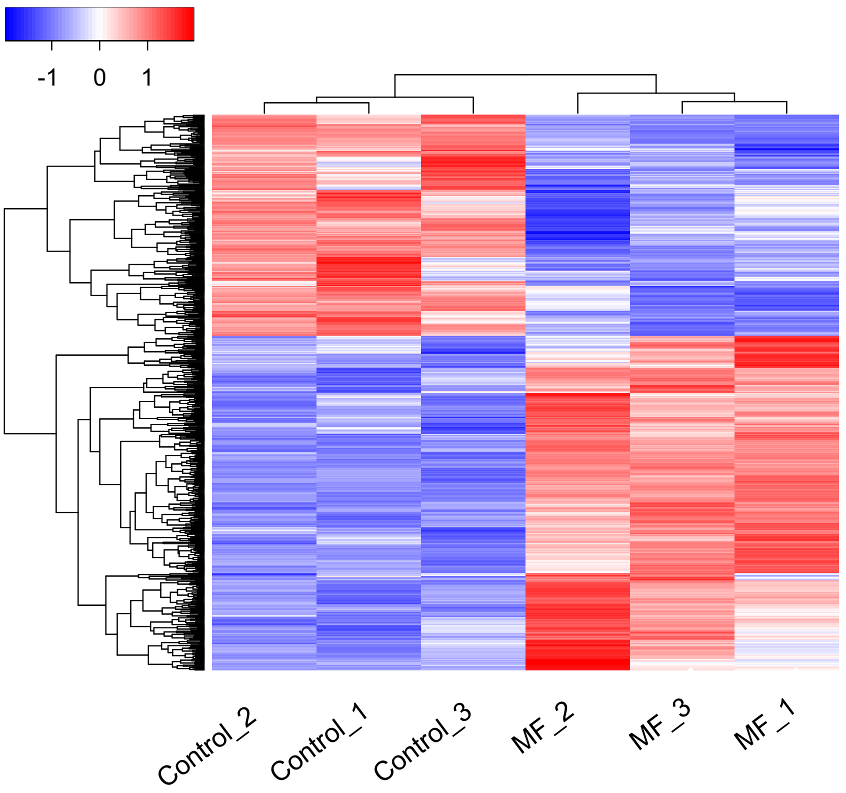


**Fig. S13.** Heatmap of the top differentially expressed genes.

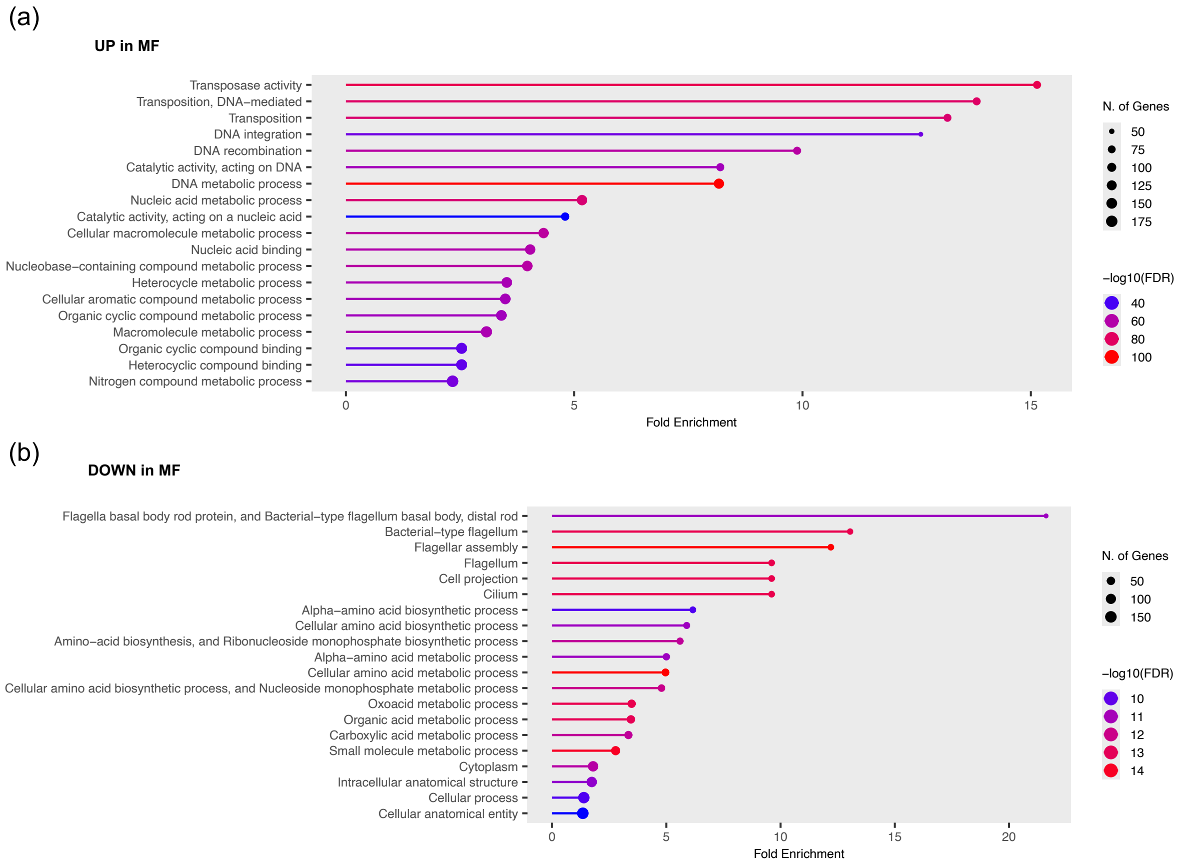


**Fig. S14.** GO enrichment analysis of top differentially expressed genes (a) upregulated and (b) downregulated in MF.

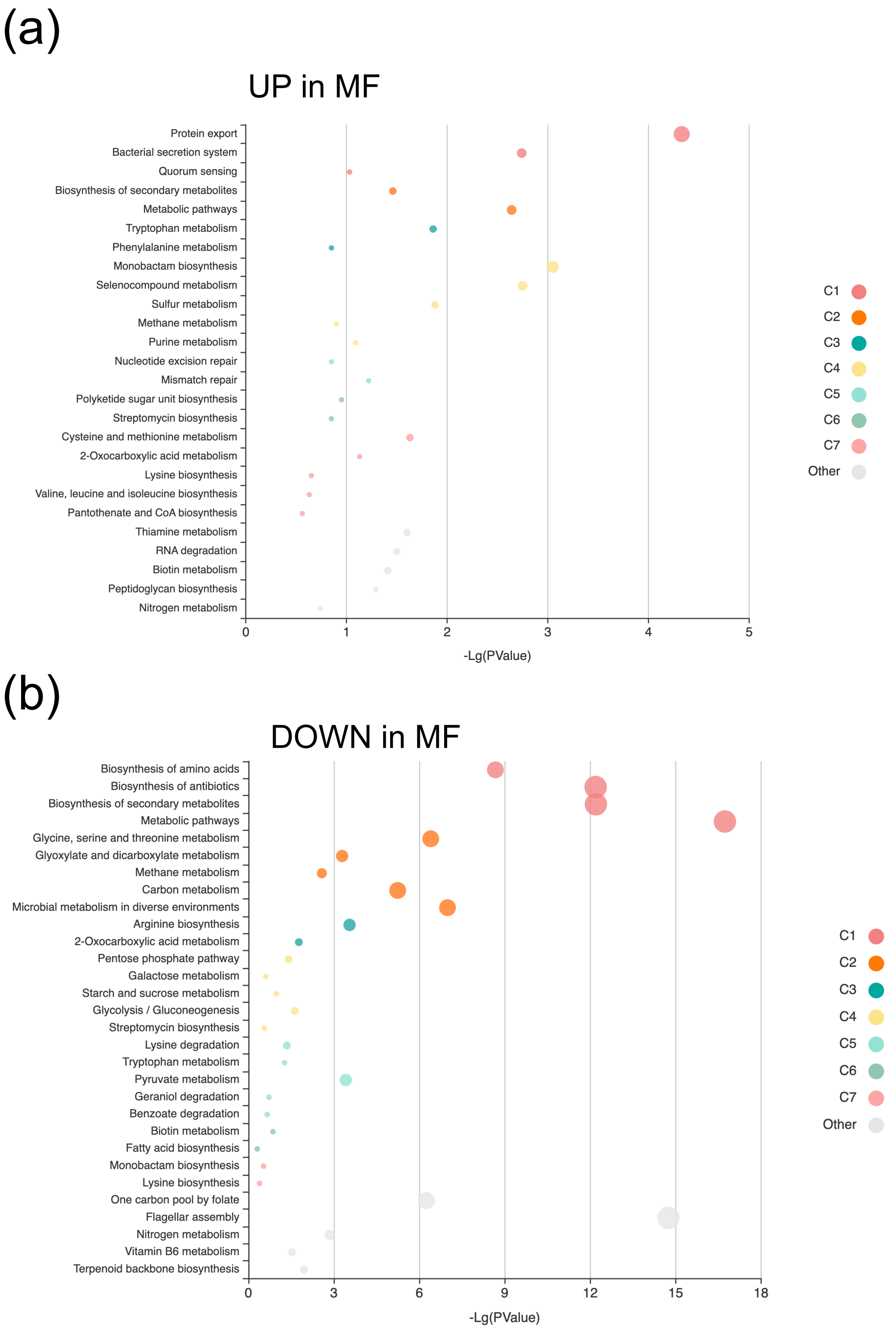


**Fig. S15.** KEGG pathway enrichment analysis of genes (a) upregulated and (b) downregulated in MF.

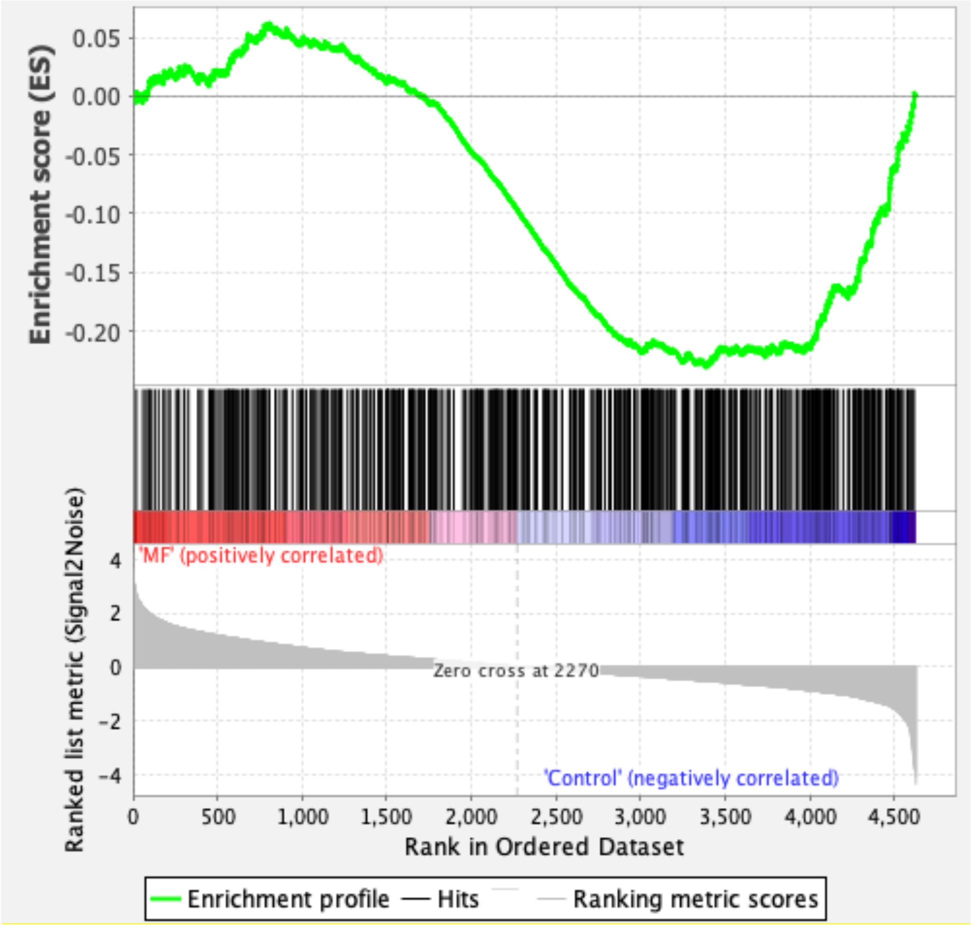


**Fig. S16.** GSEA of 481 proteins implicated in EET showing overall enrichment in control (without MF).

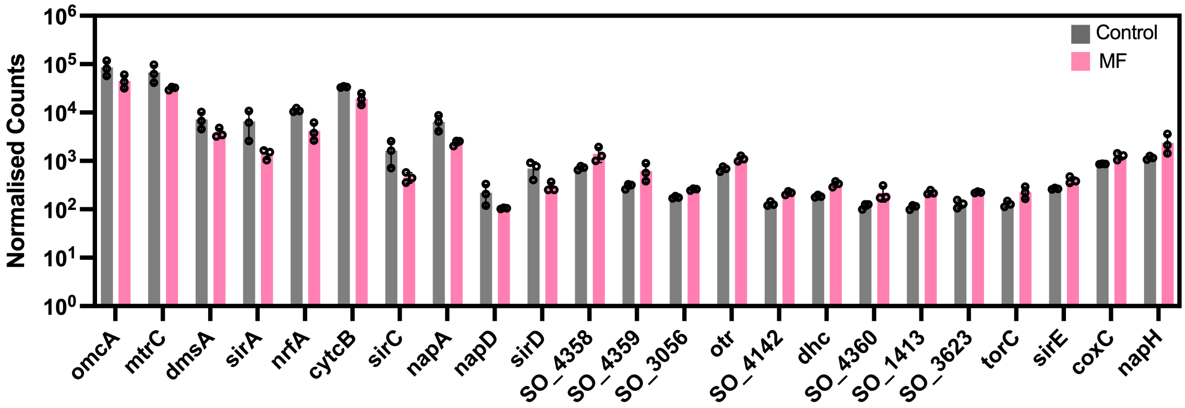


**Fig. S17.** Normalised DESeq2 counts of EET-related genes.

**Table S1.** EIS circuit-fitted values. Mean of three biological replicates shown.

| **Potential (V)** | **$R_s$ (Ω)** | **$R_B$ (Ω)** | **$R_{ct}$ (Ω)** | **$C_B$ (nF)** | **$C_{dl}$ (nF)** | **W (kσ)** |
|---|---|---|---|---|---|---|
| –0.4V [With MF] | 23.58 | 2716.66 | 52.96 | $2.06 \times 10^5$ | 2228.34 | 0.37 |
| –0.4V [Without MF] | 25.95 | 2150 | 113.35 | $3.6 \times 10^5$ | 2788.34 | 0.41 |
| 0.2V [With MF] | 21.61 | 1909 | 47.53 | $1.26 \times 10^5$ | 1515.66 | 0.35 |
| 0.2V [Without MF] | 22.94 | 1436.67 | 120.32 | $2.36 \times 10^5$ | 1800 | 0.41 |

**Table S2.** List of genes associated with EET that are differentially expressed in magnetic field.

| ***Upregulated in magnetic field*** | | | | | | | |
|---|---|---|---|---|---|---|---|
| **Gene** | **Cellular location of protein** | **$log_2$(FC)** | **P-adj** | **Product** | **GeneID** | **Old locus_tag** | **Ref. for location** |
| SO_4358 | outer-membrane (extracellular) | 0.965 | 0.0107 | dimethyl sulfoxide reductase subunit A | SO_RS20240 | SO_4358 | (*17*) |
| SO_4359 | outer-membrane | 1.033 | 0.0226 | MtrB/PioB family decaheme-associated outer membrane protein | SO_RS20245 | SO_4359 | (*17*) |
| SO_4360 | periplasm (cytochrome) | 0.902 | 0.0365 | DmsE family decaheme c-type cytochrome | SO_RS20250 | SO_4360 | (*24*) |
| SO_1413 | periplasm (cytochrome) | 1.003 | 0.0007 | cytochrome c3 family protein | SO_RS06565 | SO_1413 | (*24*) |
| dhc | periplasm (cytochrome) | 0.835 | 0.0029 | diheme cytochrome c | SO_RS20810 | SO_4485 | (*24*) |
| SO_4142 | periplasm (cytochrome) | 0.742 | 0.0124 | c-type cytochrome | SO_RS19190 | SO_4142 | (*24*) |
| otr | periplasm (cytochrome) | 0.715 | 0.0095 | tetrathionate reductase family octaheme c-type cytochrome | SO_RS19200 | SO_4144 | (*24*) |
| SO_3056 | periplasm (cytochrome) | 0.542 | 0.0448 | cytochrome c3 family protein | SO_RS14175 | SO_3056 | (*24*) |
| napH | inner membrane | 1.039 | 0.0232 | quinol dehydrogenase | SO_RS03965 | SO_0846 | (*17*) |

| | | | | | | | |
|---|---|---|---|---|---|---|---|
| | | | | ferredoxin subunit NapH | | | |
| coxC | inner membrane | 0.538 | 0.0470 | cytochrome c oxidase subunit 3 | SO_RS21410 | SO_4609 | (*17*) |
| torC | inner membrane (cytochrome) | 0.799 | 0.0470 | pentaheme c-type cytochrome TorC | SO_RS05740 | SO_1233 | (*17*, *24*) |
| SO_3623 | inner membrane (cytochrome) | 0.792 | 0.0145 | cytochrome c3 family protein | SO_RS16915 | SO_3623 | (*24*) |
| sirE | inner membrane (cyt-c maturation system) | 0.594 | 0.0400 | heme lyase NrfEFG subunit NrfE | SO_RS02285 | SO_0478 | (*17*) |
| ***Downregulated in magnetic field*** | | | | | | | |
| **Gene** | **Cellular location of protein** | **log2(FC)** | **P-adj** | **Product** | **GeneID** | **Old locus_tag** | **Ref. for location** |
| mtrC | outer membrane (extracellular) (cytochrome) | -1.081 | 0.0104 | decaheme c-type cytochrome MtrC | SO_RS08150 | SO_1778 | (*17*) |
| omcA | outer membrane (extracellular) (cytochrome) | -0.921 | 0.0493 | decaheme c-type cytochrome OmcA | SO_RS08155 | SO_1779 | (*17*) |
| dmsA | outer membrane (extracellular) | -0.907 | 0.0475 | DMSO/selenate family reductase complex A subunit | SO_RS06640 | SO_1429 | (*17*) |
| cytcB | periplasm (cytochrome) | -0.801 | 0.0140 | c-type cytochrome | SO_RS21690 | SO_4666 | (*24*) |

| | | | | | | | |
|---|---|---|---|---|---|---|---|
| sirA | periplasm (cytochrome) | -2.222 | 0.0001 | dissimilatory sulfite reductase SirA | SO_RS02290 | SO_0479 | (*24*, *25*) |
| napD | periplasm | -1.071 | 0.0372 | chaperone NapD | SO_RS03980 | SO_0849 | (*17*) |
| napA | periplasm | -1.426 | 0.0002 | nitrate reductase catalytic subunit NapA | SO_RS03975 | SO_0848 | (*17*) |
| sirC | periplasm | -1.835 | 0.0008 | 4Fe-4S dicluster domain-containing protein | SO_RS02310 | SO_0483 | (*25*) |
| nrfA | inner membrane or periplasm (cytochrome) | -1.407 | 0.0007 | ammonia-forming nitrite reductase cytochrome c552 subunit | SO_RS18515 | SO_3980 | (*17*, *24*, *25*) |
| sirD | inner membrane | -1.255 | 0.0057 | NrfD/PsrC family molybdoenzyme membrane anchor subunit | SO_RS02315 | SO_0484 | (*25*) |
| lpdA | cytoplasm | -0.529 | 0.0216 | dihydrolipoyl dehydrogenase | SO_RS02050 | SO_0426 | (*17*) |
